# ADAM10 pharmacological inhibition modifies the expression of components of the dopaminergic system

**DOI:** 10.1101/2022.05.26.493662

**Authors:** Subhamita Maitra, Bruno Vincent

**Affiliations:** Institute of Molecular Biosciences, Mahidol University, Nakhon Pathom, 73170 Thailand; Centre National de la Recherche Scientifique, 2 rue Michel Ange, 75016 Paris, France

**Keywords:** Alzheimer’s disease, neurodegeneration, cognitive deficits, sAPPα, α-secretase, dopaminergic transmission

## Abstract

Dopamine is a primary neurotransmitter associated with memory formation, emotional control, reward processing and other higher order mental functions. Altered dopamine signaling is implied in several neuropsychiatric, neurodevelopmental and neurodegenerative disorders including Alzheimer’s disease. Age-related memory decline often presents itself with spectrum of overtly behavioral responses in patients diagnosed with Alzheimer’s disease, thus suggesting that an alteration of dopaminergic transmission could account for the psychotic symptoms observed along the pathology. Since less sAPPα production due to reduced α-secretase activity is a direct contributor of compromised neuroprotection and can impart higher vulnerability to cellular insults, we explored the impact of specific inhibition of ADAM10, the main neuronal α-secretase, on dopamine system components in cultured human SH-SY5Y neuroblastoma cells. We found that dopamine receptor D4 protein levels were dose-dependently down regulated by GI254023X, but not by the ADAM17-specific inhibitor TAPI-0. We then established that GI254023X operates at a transcriptional levels. Furthermore, we showed that GI254023X treatment also significantly increased the levels of active PKA as well as the transcription of the dopamine-degrading enzymes catechol-O-methyltransferase, monoamine oxidase A and monoamine oxidase B. Altogether, our data propose that ADAM10 inhibition modulates the dopaminergic system to possibly trigger psychosis in Alzheimer’s disease.

## 1. Introduction

Mental health burden in the elderly population is growing by leaps and bounds. Frequent electrolytic imbalance, recurrent strokes and most significantly dementia are the first line contributors of aging brain hazards. Alzheimer’s disease (AD), as the first cause of dementia, is associated with severe loss of memory, lack of self -regulation and other cognitive decline. Being a complex multifactorial disorder, several processes including oxidative stress, protein aggregation, mitochondrial dysfunction and neuroinflammation are recurrent events accompanying the development of the disease.

Apart from the initial symptoms of declarative memory loss, working memory inefficiencies that gradually lead to wider range of memory loss and several domain of cognitive impairment [1], a subgroup of AD patients do exhibit psycho behavioural abnormalities characterised as behavioural and psychological symptoms of dementia (BPSD) [2,3]. The frequently noticed symptoms are apathy, aggressiveness, anxiety, depression, delusion and hallucinations [2,4,5]. Anxiety and depression are often so severe that it demands special consideration as a distinct endophenotype [6]. Importantly, AD patients with psychosis displays reduced frontal lobe function when compared to non-psychotic AD [7]. Moreover, BPSD were found to vary between late onset and early onset groups [8], are correlated to sharp cognitive decline [9] and are as heritable as the disease itself [10].

The dopaminergic system is well implied in different neuropsychiatric and neurobehavioral disorders including AD [11]. However, whether dopaminergic system alterations are genuinely involved in the course of AD is still under debate despite the existence of observations going in this direction [12,13]. At the molecular level, two members of the G_iα_-coupled D2-like family dopamine receptors D2 (DRD2) and D4 (DRD4) emerged as possibly linked to AD due to both their physiological implication in memory functions that are impaired in AD and to their previously reported alteration during the pathology. Indeed, DRD2, which is involved in memory formation in the dentate gyrus [14] and mediates cognitive flexibility in humans [15], has been shown to be reduced in AD [16] while DRD2 agonist increases cortical excitability and cholinergic transmission in AD patients [17]. As far as DRD4 is concerned, it promotes working memory performance [18-20] and an increased methylation of its promoter augments the risk of developing AD [21].

Moreover, a number of observations suggested a link between the alteration of the dopaminergic system and the amyloid pathology observed in Alzheimer’s disease. Thus, it has been shown that dopaminergic pathology and amyloid-β peptide (Aβ) deposition are closely related [22]. Moreover, it has been proposed that Aβ oligomers, through their prolonged binding to the α7 neuronal nicotinic acetylcholine receptors (α7 nAChRs), which are present on dopaminergic neurons, could alter the dopaminergic system and disrupt the mechanisms of synaptic plasticity and memory functions via an increase in calcium and an over activation of the ERK/MAPK cascade [23,24]. However, whether modulating the production of other β-amyloid precursor protein (βAPP)-derived metabolite through the regulation of secretases could interfere with components of the dopaminergic system still remains elusive. Such a possibility is actually supported by the fact that the treatment of rat hippocampal organotypic slices with 1nM of the βAPP α-secretase-derived sAPPα metabolite for 24 hours induces a 1.29-fold increase in DRD2 expression [25].

We here undertook to investigate the impact of a direct and specific inhibition of ADAM10 on the levels of DRD2, DRD4 and several other components of the dopaminergic systems in the human neuroblastoma SH-SY5Y cell line. Importantly, because GI254023X possesses a 100 times greater inhibitory capacity on ADAM10 compared to ADAM17 recombinant proteases [26,27], since it is established that ADAM10 is the main physiological α-secretase activity in neurons [28] and because this inhibitor completely abrogates IGF-1-induced sAPPα production in SH-SY5Y cells at concentrations similar to those used in the present study [29], we can reasonably consider this inhibitor to be specific for ADAM10 under our experimental conditions.

Our data show that GI254023X, but not the ADAM17-specific inhibitor TAPI-0, dose-dependently decreased DRD4 expression without altering DRD2 levels. Moreover, GI254023X also increased the immunoreactivity of the catalytic subunit α of PKA as well as the mRNA levels of the dopamine-degrading catechol-O-methyltransferase (COMT), monoamine oxidase A (MAOA) and monoamine oxidase B (MAOB) enzymes. Altogether, our results established that ADAM10 specific inhibition perturbs the dopaminergic system and might be part of the molecular bridge linking AD pathology with its associated psychotic symptoms.

## 2. Materials and Methods

### 2.1. Antibodies, reagents and cell lines

Polyclonal anti-ADAM10 (AB19026) and anti-ADAM17 (AB19027) antibodies were purchased from Millipore (Bedford, MA, USA). Polyclonal anti-βAPP (A8717), monoclonal anti-β-actin (A2228), dimethyl sulfoxide (DMSO) and sodium bicarbonate were from Sigma (St Louis. MO. USA). Monoclonal anti-DRD2 (sc-5303), anti-DRD4 (sc-136169) and anti-GAPDH (sc-32233) antibodies were from Santa Cruz (Santa Cruz, CA, USA). The monoclonal anti-β-amyloid antibody (DE2B4), which was used to specifically detect sAPPα, was from IBL (Minneapolis, MN, USA). The polyclonal antibody specifically recognizing the catalytic subunit α of PKA (4782), the goat anti-mouse (polyclonal 7076) and goat anti-rabbit (polyclonal 7074) peroxidase-conjugated secondary antibodies were from Cell Signaling (Beverly, MA. USA). DMEM complete medium, foetal bovine serum (FBS) and penicillin-streptomycin mix were from Invitrogen (Carlsbad, CA, USA). ECL and ammonium persulphate (APS) were from GE Healthcare (Pisataway, NJ, USA). Tris buffer, glycine and sodium dodecyl sulphate (SDS) were from Amresco (Solon, CA, USA). Skim milk powder was from Bio Basic (Singapore). Human SH-SY5Y neuroblastoma cells (gift from Dr. Narisorn Kitiyanant, Mahidol University) were grown at 37°C, 5% CO_2_ in high glucose-DMEM supplemented with 10% FBS, penicillin (100U/ml) and streptomycin (50mg/ml).

### 2.2. ADAM10 and ADAM17 specific inhibitions with GI254023X and TAPI-0 respectively

The stock solutions of GI254023X (Sigma) and TAPI-0 (Calbiochem, San Diego, CA, USA) were prepared at 10mM in 100% DMSO from which different intermediate concentrations were prepared in 10% DMSO and kept frozen. The cells were treated with the respective intermediate solutions for 15 hours in complete media. The final concentration of DMSO in each treated well was 0.1%. Therefore, control cells were always vehicle-treated with 0.1% DMSO.

### 2.3. Western blot analysis

At the end of the 15 hours treatments, 30-40μg of proteins were separated by SDS-polyacrylamide gel electrophoresis on 10% (ADAM10 and ADAM17) or 12% (DRD2, DRD4 and PKA) Tris/glycine gels and run at 100 volts for 2-2.5 hours. Proteins were then transferred onto a nitrocellulose membrane for 60-90 minutes at 90 volts. After the effectiveness of protein transfer was checked with Ponceau red staining, nitrocelluloses were incubated in 5% non-fat milk blocking solution for 30 min. Membranes were then incubated with primary antibodies directed against βAPP (1/4000 dilution), ADAM10 (1/500 dilution), ADAM17 (1/1000 dilution), DRD2 (1/500 dilution), DRD4 (1/500 dilution) or PKA Cα (1/1000 dilution) on a platform shaker overnight at 4°C. On the next day, after 3 washes with PBST (PBS containing 0.05% Tween 20), membranes were incubated with HRP-conjugated anti-rabbit (βAPP, ADAM10, ADAM17, PKA Cα) or anti-mouse (D2R, D4R) secondary antibodies (1/3000) for 2h and rinsed 3 times with PBST. Immunoreactivities were processed using ECL and signals were detected using the Azure C400 Gel Imaging System. Quantification of band densities was performed using the Image J Analyzer software (http://imagej.nih.gov/ij/). All membranes were subsequently rebloted with GAPDH (1/7500) or β-actin (1/5000) and the HRP-conjugated anti-mouse secondary antibody. GAPDH and β-actin immunoreactivities were used as internal standard to normalize the data.

### 2.4. sAPPα secretion and measurement

SH-SY5Y cells were cultured in 35mm-dishes until they reached 80% confluence. Cells were incubated without (control) or with GI254023X or TAPI-0 (100nM, 1μM, 10μM) for 15 hours at 37^0^C in 1ml of serum-free DMEM. TCA (10%) precipitation was then performed from the whole media (1ml) and the entire protein precipitate was submitted to western blot analysis on a 10% SDS-PAGE using the primary antibody DE2B4 directed against sAPPα (1/500 dilution) and the secondary HRP-conjugated anti-mouse antibody and processed as described above.

### 2.5. Real-time quantitative polymerase chain reaction (q-PCR)

Following treatments for 15 hours in complete media, total RNA was extracted and purified with the PureLink RNA mini kit (Ambion, Life Technologies, Austin, TX, USA). Real-time PCR was performed with 100ng of total RNA using the QuantiFast SYBR Green RT-PCR kit (Qiagen, Singapore) detector system (ABI Quant studio 5) and the SYBR Green detection protocol. The 2x QuantiFast SYBR Green RT-PCR master mix, QuantiFast RT mix, QuantiTectPrimer Assay and template RNA were mixed and the reaction volume was adjusted to 25μl using RNase-free water. The specific primers were designed and purchased from Qiagen. Each primer is a 10x QuantiTect Primer Assay containing a mix of forward and reverse primers for specific targets: Hs_DRD2_1 (QT00012558), Hs_DRD4_1 (QT00204316), Hs-DBH_1 (QT00047194), Hs_DDC_1 (QT00046774), Hs_COMT_2 (QT01890756), Hs_MAOA_1(QT00040411), Hs_MAOB_1 (QT-00009870) and Hs_GAPDH_1_SG (QT00079247, human GAPDH).

### 2.6. Cell viability assay

Cells were seeded in 96-well polystyrene-coated tissue culture plate (corning) over night. Proper attachment and confluence of the cells were confirmed by checking under microscope. Media was removed and cells were treated with inhibitors at various concentrations in quadruplicate for different time points with control cells being treated with vehicle (0.1% DMSO). Following treatment for 15 hours, (3-(4,5-dimethylthiazol-2-yl)-2,5-diphenyltetrazolium bromide (MTT) (0.5mg/ml) was added for 1.5 hours. Formazan crystals formation was then checked under microscope and the media was replaced by DMSO (100%) until purple crystals have dissolved. Absorbance was measured at 595nm.

### 2.7. Statistical analysis

Statistical analyses were performed with the Prism software (GraphPad, San Diego, USA) using the unpaired t test for pair-wise comparisons. All values were the means ± SE with *p* values equal to or less than 0.05 being considered significant.

## 3. Results

### 3.1. GI254023X impairs sAPPα production and decreases DRD4 protein levels

We first examined the effect of 15 hour-treatments of cultured SH-SY5Y cells with the specific ADAM10 inhibitor GI254023X at three different doses (0.1, 1 and 10μM) on the production of sAPPα (the βAPP-derived metabolite engendered by ADAM10) as well as on the protein levels of βAPP (ADAM10 substrate) and ADAM10 itself. The results showed a dose-dependent decrease in sAPPα secretion by GI254023X with about 75% inhibition at 10μM (Figure 1A and E) with no modification of βAPP (Figure 1A and F) and ADAM10 (Figure 1B and G) immunoreactivities whatever the dose considered. These observations are in good agreement with our previous data having established that 10μM of GI254023X only impairs sAPPα production without modifying βAPP and ADAM10 levels in the same cell line [30]. We then wanted to determine, as part of our initial objective, whether GI254023X could modify the pattern of DRD2 and DRD4 protein levels under the same experimental conditions. Although no changes were observed regarding DRD2 protein levels following GI254023X treatment at all concentrations (Figure 1C and H), DRD4 immunoreactivity was gradually and significantly diminished by GI254023X with an inhibition profile quite similar to the one observed for sAPPα (Figure 1D and I).

**Figure 1.**
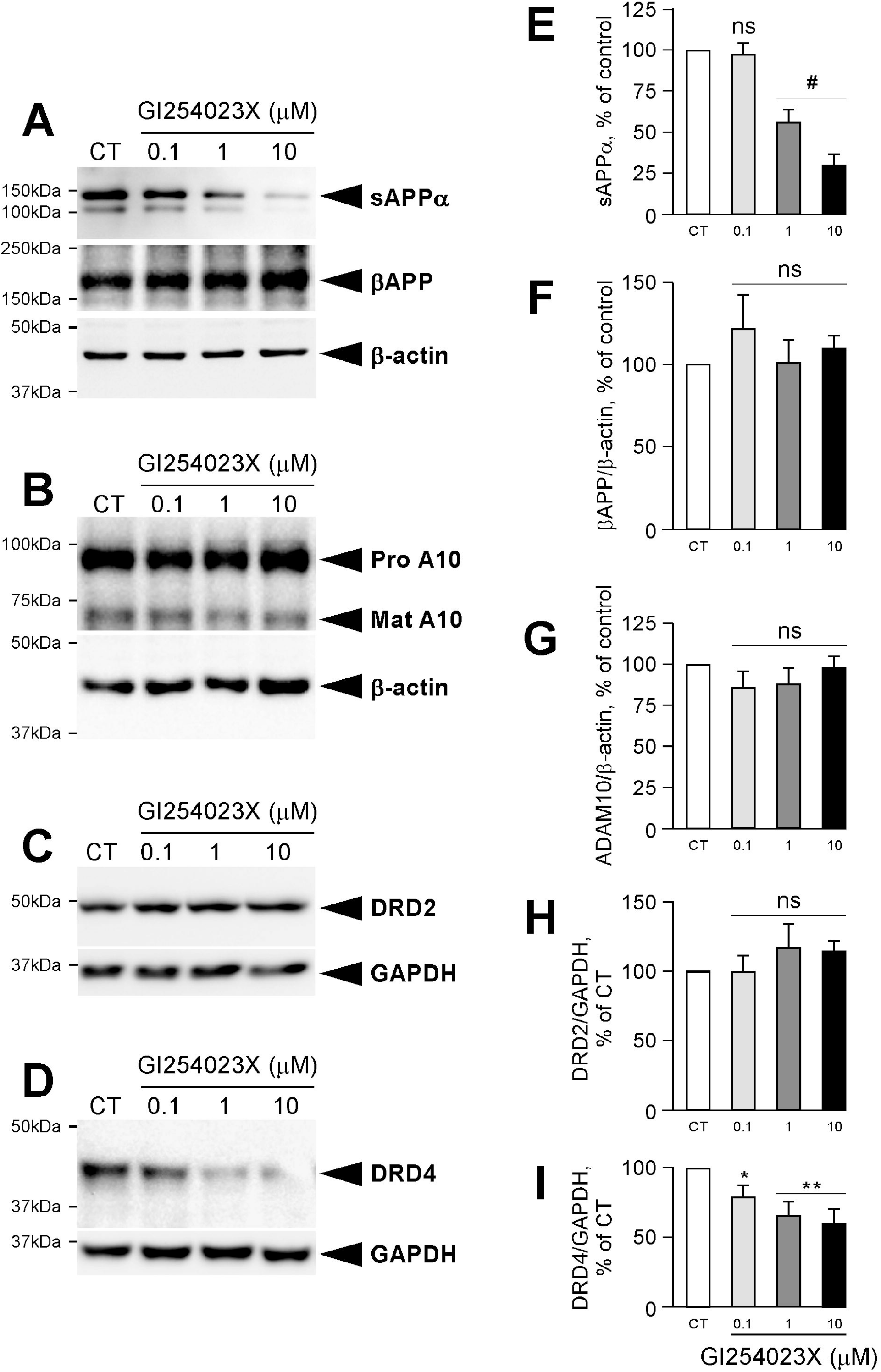
Effect of GI254023X on sAPPα production and βAPP, ADAM10, DRD2 and DRD4 protein levels in human SH-SY5Y cells. Left panels illustrate representative gels of western blot analysis of sAPPα production in media (A) as well as βAPP (A), ADAM10 (B), DRD2 (C) and DRD4 (D) and their respective GAPDH/β-actin in lysates following treatment of cultured SH-SY5Y cells without (CT) or with the indicated concentrations of GI254023X for 15 hours. (B) Right panels show the statistical analysis of the data for sAPPα (E), βAPP (F), ADAM10 (G), DRD2 (H) and DRD4 (I). Bars correspond to the densitometric analyses (βAPP and ADAM10 being normalized with β-actin, DRD2 and DRD4 being normalized with GAPDH), are expressed as a percentage of control and are the means ± SE of 11 (sAPPα), 9 (βAPP, DRD2 and DRD4) or 6 (ADAM10) independent determinations. *p≤0.05; **p≤0.01; #p≤0.0001; ns, non-statistically different (p>0.05).

### 3.2. Specific inhibition of ADAM17 does not modify DRD2 and DRD4 immunoreactivities

We then examined the impact of the specific inhibition of ADAM17, another α-secretase activity barely involved in the constitutive metabolism of βAPP, on the same parameters. Firstly, treatments with the ADAM17 specific inhibitor TAPI-0 did not show any decrease in the production of sAPPα (Figure 2A and E) and in the levels of βAPP (Figure 2A and F). The absence of participation of this enzyme in the α-secretase cleavage of βAPP under our experimental conditions is in accordance with the fact that ADAM17 is principally involved in the PKC-regulated α-secretase cleavage of βAPP [31]. Secondly, beyond the fact that TAPI-0 did not change ADAM17 immunoreactivity (Figure 2B and G), it also failed to show an effect on both DRD2 (Figure 2C and H) and DRD4 (Figure 2D and I) protein levels.

**Figure 2.**
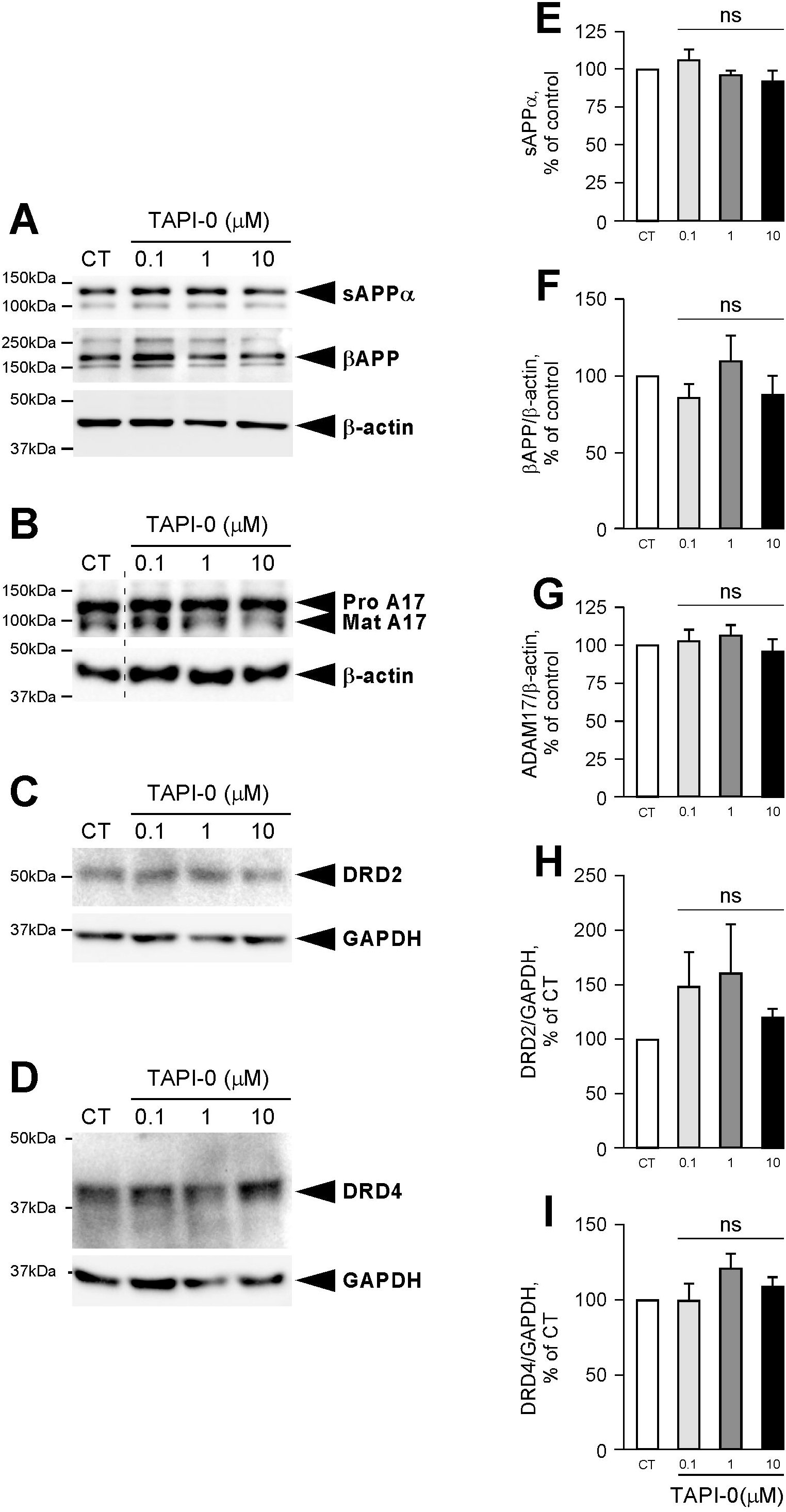
Effect of TAPI-0 on sAPPα production and βAPP, ADAM17, DRD2 and DRD4 protein levels in human SH-SY5Y cells. Left panels illustrate representative gels of western blot analysis of sAPPα production in media (A) as well as βAPP (A), ADAM17 (B), DRD2 (C) and DRD4 (D) and their respective GAPDH/β-actin in lysates following treatment of cultured SH-SY5Y cells without (CT) or with the indicated concentrations of TAPI-0 for 15 hours. Right panels show the statistical analysis of the data for sAPPα (E), βAPP (F), ADAM17 (G), DRD2 (H) and DRD4 (I). Bars correspond to the densitometric analyses (βAPP and ADAM17 being normalized with β-actin, DRD2 and DRD4 being normalized with GAPDH), are expressed as a percentage of control and are the means ± SE of 6 (βAPP and ADAM17), 5 (sAPPα) or 3 (DRD2 and DRD4) independent determinations. ns, non-statistically different (p>0.05). The hatched black line in (B) indicates a splicing of the original gels.

### 3.3. GI254023X and TAPI-0 treatments do not trigger cellular toxicity

At this stage, it was important to rule out the existence of a possible cellular toxicity induced by a prolonged exposure to these inhibitors as performed under our experimental conditions. With the aim of ruling out such a hypothesis, we measured the survival rate of SH-SY5Y cells following a 15 hour treatment without (control) or with GI254023X or TAPI-0 at 0.1, 1 and 10μM concentrations by means of the MTT test. The results showed that no notable changes were observed whatever the concentrations considered when compared to the non-treated cells (Figure 3), obviously indicating that GI254023X and TAPI-0 were not conveying toxicity under our experimental conditions as further confirmed by the absence of cell morphology differences between treatment conditions.

**Figure 3.**
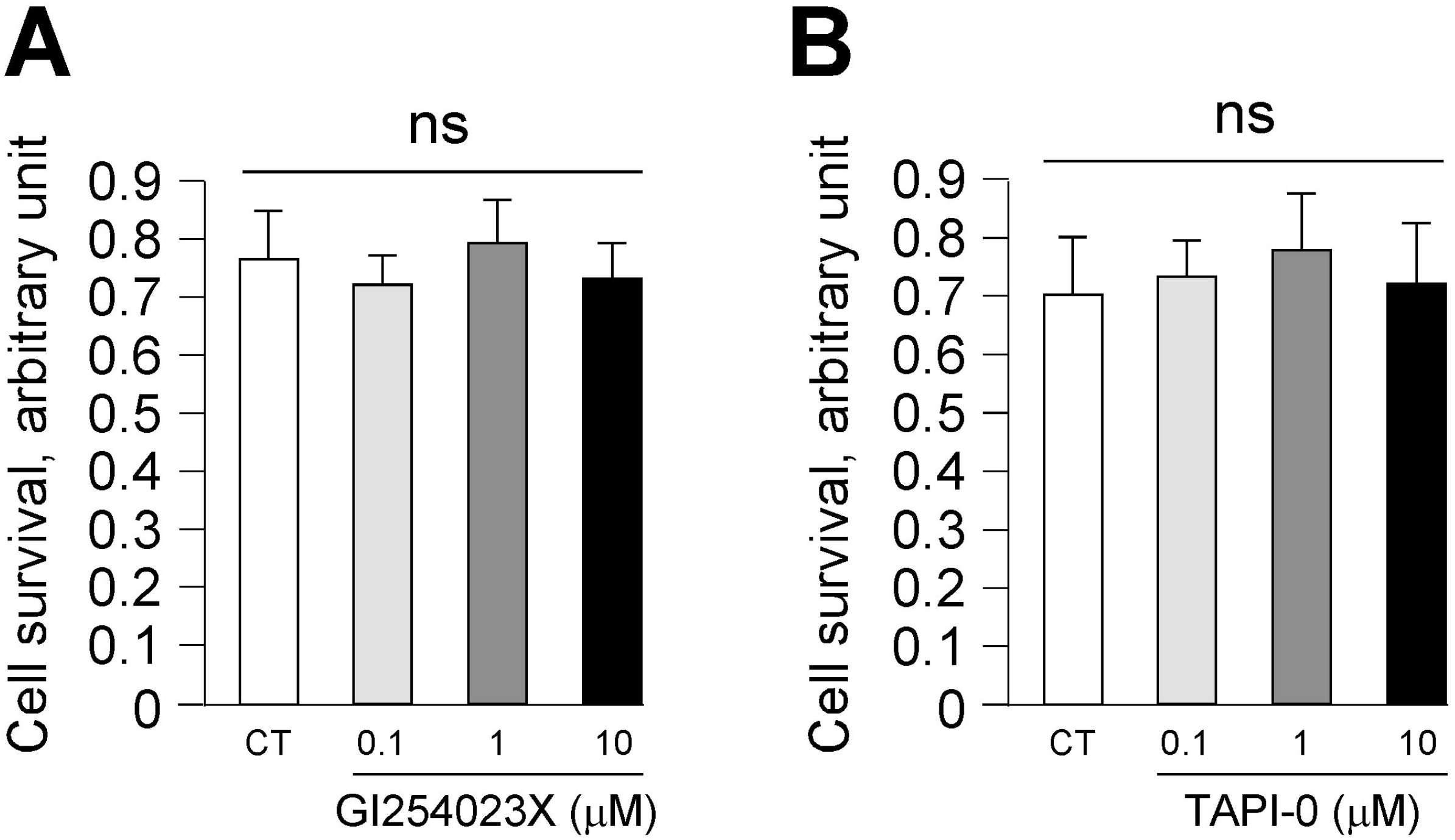
Effect of GI254023X and TAPI-0 on cellular toxicity in SH-SY5Y cells. Cells were incubated without (control, white bar) or with the indicated concentrations of GI254023X (A) or TAPI-0 (B) for 15 hours and cell viability was determined with the MTT assay. The results are expressed as the mean of 14 (A) or 11 (B) independent determinations. ns, non-statistically different (p>0.05).

### 3.4. ADAM10 specific inhibition decreases DRD4 and increases COMT, MAOA and MAOB at a transcriptional level

In order to establish whether the lessening of DRD4 protein levels mediated by ADAM10 specific inhibition was reflecting a down regulation of its transcription, we undertook to measure the impact of the treatment with GI254023X at the more efficient concentration (10μM) on the mRNA levels of DRD2 and DRD4 by RT-PCR.

Additionally and in order to broaden our field of investigation, we also examined the transcriptional patterns of several other important components of the dopaminergic system. After the careful checking of which of such genes were expressed in SH-SY5Y cells (https://www.proteinatlas.org/), we selected dopamine decarboxylase (DDC), dopamine-β-hydroxylase (DBH), monoamine oxidase (MAO) A and B as well as catechol-O-methyltransferase (COMT) for analysis.

The results first showed that, as observed at the protein level, GI254023X does not alter the amounts of DRD2 mRNA while significantly decreasing DRD4 mRNA levels (Figure 4A and B), thereby indicating that ADAM10 inhibition diminishes DRD4 by triggering some mechanisms leading to a modulation of its transcription. Regarding the regulation of some other elements playing important roles in dopaminergic transmission, we first did not evidence any effect of GI254023X treatment on the mRNA levels of the DDC and DBH enzymes (Figure 4C and D), thus indicating that ADAM10 inhibition most likely neither affects the synthesis of dopamine L-DOPA nor the conversion of dopamine to norepinephrine. Interestingly, however, we were able to note a significant stimulating effect of treatment with GI254023X on the transcription of three enzymes responsible for dopamine degradation, namely COMT, MAOA and MAOB (Figure 4E-G).

**Figure 4.**
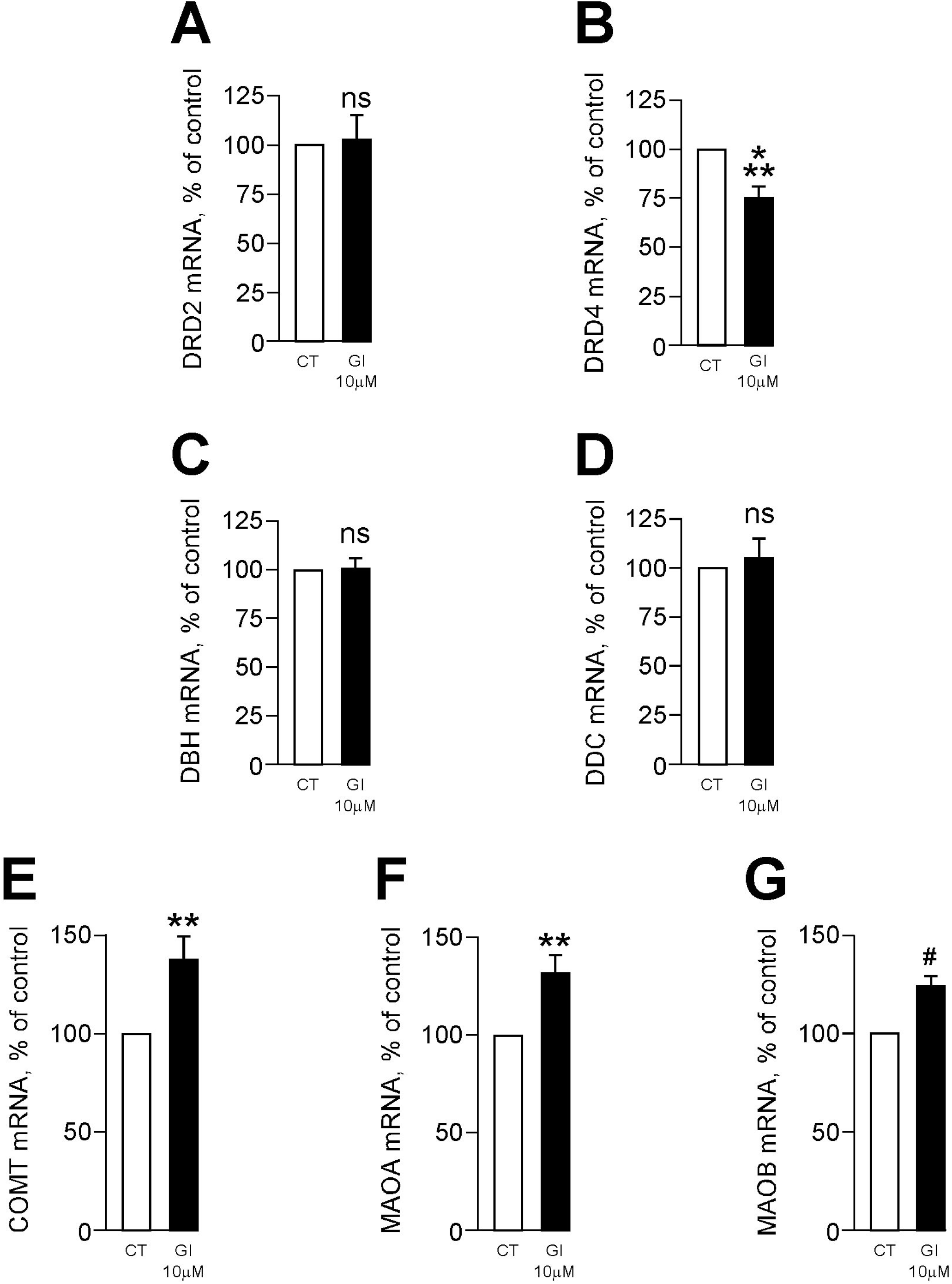
Effect of GI254023X on the mRNA levels of components of the dopaminergic system. Following treatment of SH-SY5Y cells with GI254023X (10μM) for 15 hours, DRD2 (A), DRD4 (B), DBH (C), DDC (D), COMT (E), MAOA (F) and MAOB (G) mRNA levels (normalized with GAPDH) were measured by real-time qPCR. Bars in black are expressed as a percentage of control (white bar) and are the means ± SE of 12 to 18 independent determinations. ** p≤0.01; ***p≤0.001; #p≤0.0001; ns, non-statistically different (p>0.05).

### 3.5. GI254023X triggers an increase in active PKA Cα immunoreactivity

Finally, in order to correlate the observed GI254023X-mediated decrease in DRD4 expression with some DRD4-dependent downstream signaling events, we examined the effect of a loss of ADAM10 activity on the levels of the catalytic subunit α of PKA that is down regulated by this receptor via the inhibition of adenylate cyclase and the subsequent decrease in cAMP production. The results showed that GI254023X significantly and dose-dependently increased PKA Cα protein immunoreativity (Figure 5).

**Figure 5.**
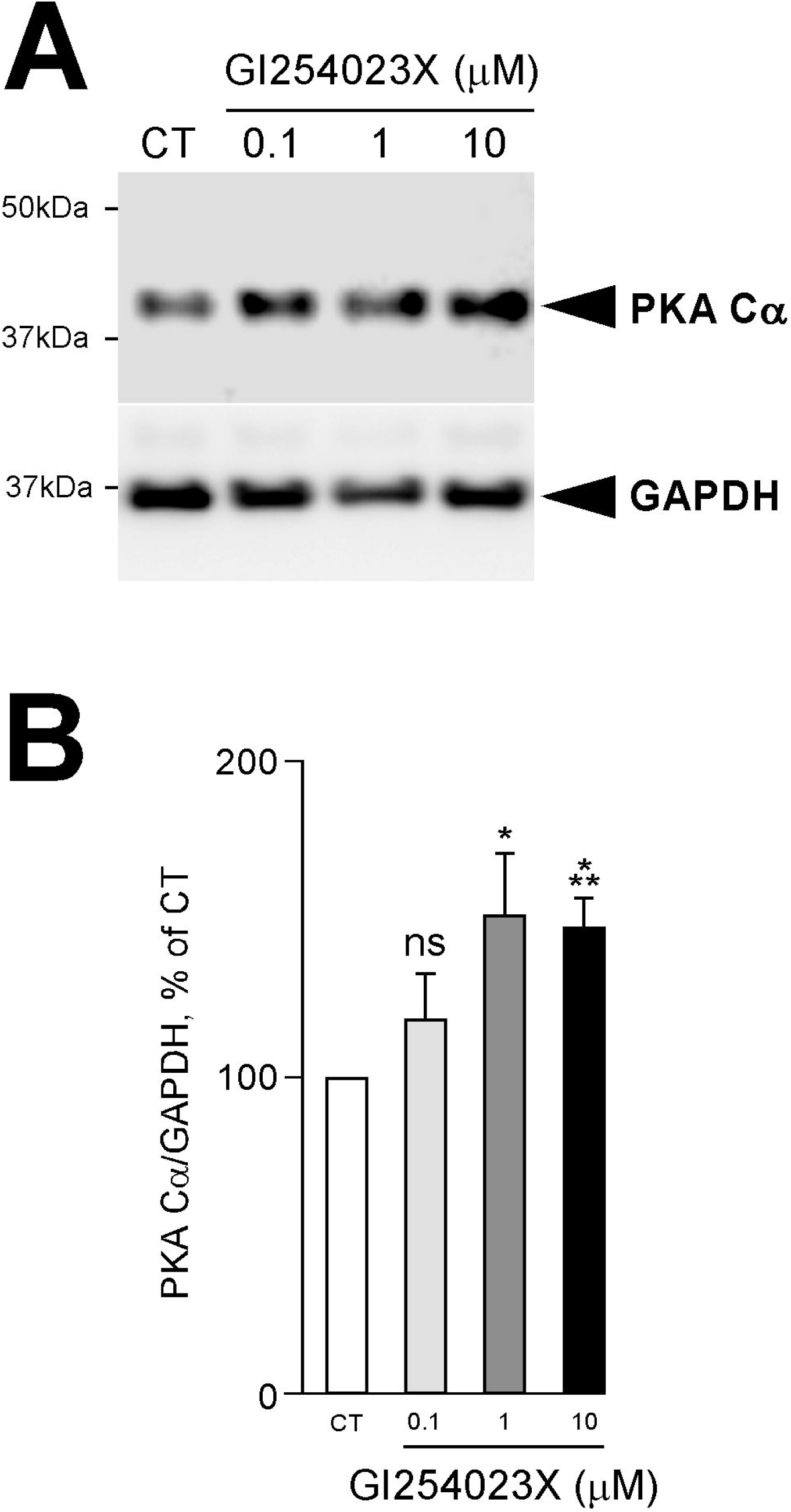
Effect of GI254023X on the levels of the catalytic unit α of PKA. (A) illustrates a representative gels of western blot analysis of the catalytic unit α of PKA and its corresponding GAPDH in lysates following treatment of cultured SH-SY5Y cells without (CT) or with the indicated concentrations of GI254023X for 15 hours. (B) shows the statistical analysis of the data. Bars correspond to the densitometric analyses (PKA Cα being normalized with GAPDH), are expressed as a percentage of control and are the means ± SE of 9 independent determinations. * p≤0.05; *** p≤0.001; ns, non-statistically different (p>0.05).

## 4. Discussion

Being a multifactorial complex disorder, the exact aetiology of AD is very difficult to pinpoint as multiple interactive pathways are involved. Nevertheless, it is nowadays largely consensual that a deregulation of the processing of βAPP is the trigger of a cascade of events involving synergistically toxic Aβ species and hyperphosphorylated tau [32]. Since it takes place at the very early stages of the disease, interfering with the differential processing of βAPP by secretases has long been considered as a pertinent therapeutic way to combat AD pathology. On the one hand, the β-secretase BACE1, as the limiting factor of Aβ production, is a first line leakage through which the pathological cascades start penetrating and is therefore widely considered a therapeutic target of choice [33], although its involvement in the cleavage of other substrates supporting important physiological functions brings a downside to this postulate [34]. On the other hand, the non-amyloidogenic processing of βAPP by α-secretase prevents Aβ formation and promotes, through the production of the sAPPα metabolite, neuroprotection, neurotrophism, memory performance, synaptic plasticity and neurogenesis [35]. Considering that α- and β-secretase compete for βAPP processing, that β-secretase activity is increased both during normal aging [36] and in AD [37], and given the fact that reduced α-secretase and deficits in sAPPα levels were observed in AD patients [38], restoration of α-secretase activity can be seen as a vital aspect for AD therapeutic intervention [39]. In this context, ADAM10, the main neuronal physiological α-secretase [28], is centre stage to the non-amyloidogenic metabolism of βAPP that can potentially cut short the early deleterious events leading to the disease.

Importantly, a certain number of previously reported observations support a probable implication of dopamine (DA) in AD. Firstly, the fact that lower levels of dopamine was reported in AD [13] together with the observation that one third of AD patients present extrapyramidal signs [40], strongly suggest that dopaminergic neurons undergo degeneration during the time course of the disease. Secondly, neurons from the nigrostriatal pathway are not only tightly associated with AD-related pathologic changes such as neurofibrillary tangles (NFT) and senile plaques [41-44], but also show decreased DA content. [45]. Thirdly, the mesocorticolimbic system, another major dopaminergic pathway involved in higher order mental functions, motivation, mood and emotional valence, was also found to undergo atrophy [46]. Finally, additional works carried out in transgenic mouse models of AD have convincingly strengthen the link between the disease and DA system failure as shown by the fact that dopamine neuronal loss contributes to memory dysfunction in Tg2576 mice [47] while, conversely, the restoration of DA transmission alleviates memory and learning in TgCNRD8 [48] and in 3xTg-AD mice [49].

A role for some βAPP-derived metabolites in modulating the dopaminergic system have been brought up in recent years. Thus, intracerebral injection of Aβ_42_ oligomers (produced from the amyloidogenic β/γ processing of βAPP) decrease cortical dopamine levels *in vivo* in mice [50] while sAPPα (issued from the non amyloidogenic α processing of βAPP) increases by 30% DRD2 mRNA levels in rat hippocampal slice cultures [25]. In this context, we aimed to investigate the effect of ADAM10 inhibition on components of the dopaminergic system endogenously expressed in the human neuroblastoma SH-SY5Y cell line.

We first showed that the specific inhibition of ADAM10 by GI254023X, which as expected leads to a dose-dependent lessening of sAPPα production, has different impacts on the protein and mRNA levels of DRD2 and DRD4, two members of the D2-like family of dopamine receptors that are both implicated in memory functions and altered in AD. Thus, GI254023X specifically and dose-dependently reduced DRD4 immunoreactivity without modifying DRD2 levels. At the same time, quantitative PCR analysis of the impact of the GI254023X concentration producing maximum effect on DRD4 protein levels (10μM) went in the same direction with a significant decrease in DRD4 mRNAs and no effect on DRD2. Importantly, our observation that the specific inhibition of ADAM17 by TAPI-0 had no impact on both sAPPα production and DRD4 levels indicated, beyond the fact that the pharmacological inhibitions of two closely related protease from the same family convey distinct effects, that the reduction of DRD4 by GI254023X could putatively be due to a decrease in sAPPα production.

These results may appear contradictory to the finding that treatment with exogenous sAPPα triggers a 1.29-fold up-regulation of DRD2 mRNA levels [25]. This discrepancy could be explained by the implementation of different approaches aimed at modulating the levels of circulating sAPPα (pharmacological inhibition of α-secretase versus exogenous addition of sAPPα; 15h versus 24h treatments) as well as the models used (human cultured neuroblastoma cells versus rat hippocampal slice cultures). Nonetheless, additional experiments would certainly help to precisely decipher the global implication of the non amyloidogenic processing of βAPP on the expression profiles of dopaminergic receptors.

Our additional observation that GI254023X increases the availability of the catalytic subunit α of PKA (PKA Cα) added further evidenced the impact of ADAM10 inhibition on DRD4 since these receptors inhibit this kinase via the repression of adenylate cyclase activity and the subsequent reduction in cAMP production. In the context of AD pathogenesis, it is interesting to note that PKA over activation has been shown to increase tau phosphorylation and oxidative stress [51] and to shorten neuronal dendrites in drosophila [52] and rats [53]. Moreover, it has been evidenced that hippocampal PKA over activation disrupts recognition and spatial memory [54]. It is also important to note here that the observed increase in PKA Cα levels induced by GI254023X could result, independently of the inhibition of DRD4 expression, from a decrease in the GABAb receptor-dependent signaling pathway. Indeed, sAPPα having been clearly established as a ligand for GABAbR1a [55], it is expected that a decrease in its production will lead to an increase in PKA activation following the reduction in the inhibitory tone of these receptors on adenylate cyclase [56].

Beyond the effects observed on dopaminergic receptors, we have also established that GI254023X, at the concentration of 10μM that drastically impairs sAPPα production, also increased the mRNA levels of the three principal dopamine-degrading enzymes COMT, MAOA and MAOB. It has to be underline here that these three enzymes have been genetically and/or biochemically associated with AD. Thus, the COMT GG genotype (Val/Val) at position 158, where there is a single nucleotide polymorphism (SNP) (G/A; Val/Met), with the Val allele exhibiting a 3- to 4-fold increase in enzyme activity compared to Met, is synergistically associated with the APOE ε4 allele in increasing the risk of AD and mild cognitive impairment (MCI) [57]. In addition, several works have pinpointed some links between MAOA and AD. Firstly, the levels of MAOA mRNA are increased in the frontal cortex of AD patients [58]. Secondly, an association of MAOA polymorphism (GT dinucleotide repeat in MAOA allele 113 gene) with AD has been evidenced [59]. Finally, a promoter polymorphism (high number of tandem repeats) in the MAOA gene was associated with higher MAOA gene expression and activity in AD patients [60]. Regarding MAOB, it has been shown that its activity was increased up to three-fold in the temporal, parietal and frontal cortices of AD cases when compared with controls [61]. Moreover, it has been recently evidenced that MAOB levels were higher in the frontal cortex, hippocampus CA1 and entorhinal cortex of AD brain than in controls, that the enzyme is a γ-secretase-associated protein and that intraneuronal Aβ_42_ levels correlated with MAOB levels [62]. Importantly, this molecular and genetic evidence has recently been reinforced by bioinformatics studies using microarray datasets that have detected higher expression of MAOA, MAOB and COMT in the brain of AD patients when compared to control individuals [63-65] and modified COMT expression in the hippocampus of AD patients [66]. Furthermore, a meta analytic review has also reported a significant increase in hippocampal MAOB expression in AD [67]. Altogether and in line with our results, these data support the idea that during the time course of AD, an excess in MAOA, MAOB and COMT activities could contribute to an exacerbated dopamine metabolism, thus designating these enzymes as a possible therapeutic targets for the treatment of the disease [68].

## 5. Conclusions

Altogether, our results established that ADAM10 specific inhibition by GI254023X triggers an overall dopaminergic tonus breakdown manifested on one hand by a decrease in DRD4 expression leading to an augmentation of the levels of the catalytic unit α of PKA and on the other hand to an increase in the levels of the dopamine-degrading enzymes that altogether could account for some of the cognitive and behavioral impairments observed in the course of AD (Figure 6).

**Figure 6.**
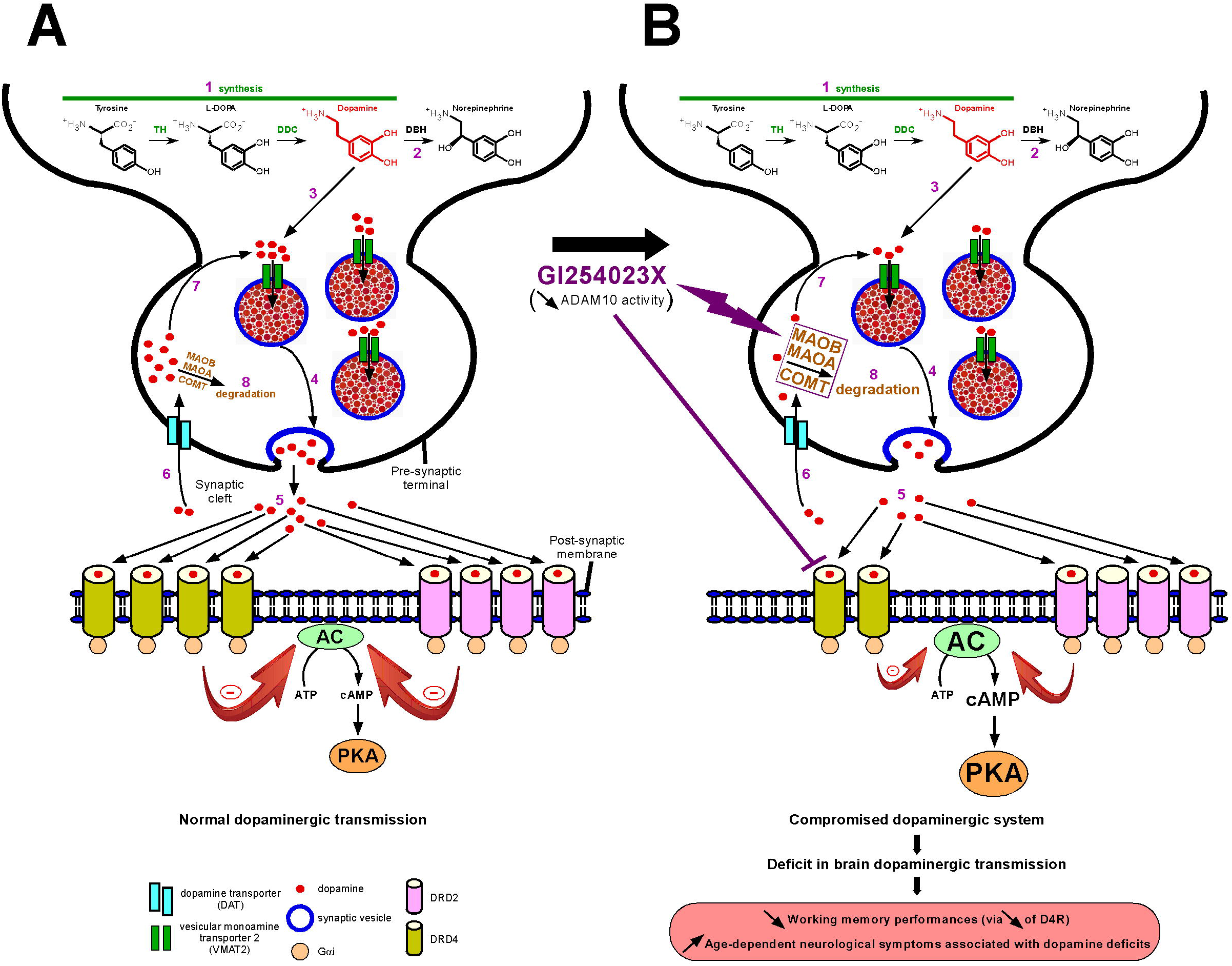
Schematic representation of the effects of GI254023X on the dopaminergic system, their consequences on the dopaminergic transmission and their possible impact in promoting AD pathogenesis. (A) Illustration of the dopamine system under normal conditions. Dopamine (DA) is first synthetized in a two-step mechanism via the conversion of tyrosine into L-DOPA by tyrosine hydroxylase (TH) and the subsequent transformation of L-DOPA into dopamine by DOPA decarboxylase (DDC) in the presynaptic terminal of dopaminergic neurons (1). Dopamine can then either be converted into norepinephrine by dopamine-β-hydroxylase (DBH) (2) or sequestered into synaptic vesicles by the vesicular monoamine transporter 2 (VMAT2) (3) from where it is released in the synaptic cleft (4). After the binding of DA to its receptors located at the postsynaptic membrane (here only DRD2 and DRD4 are depicted) and the perpetuation of signaling transduction (5), it is recycled (via recapture by the dopamine transporter (DAT)) to the cytoplasm (6) where it can either reincorporate synaptic vesicles through VMAT2 (7) or be degraded by monoamine oxidase A and B (MAOA and MAOB) and catechol-O-methyltransferase (COMT) (8). (B) Illustration of the changes observed in the dopaminergic system following specific ADAM10 inhibition. GI254023X treatment triggers two main effects. Firstly, it selectively reduces the expression/transcription of DRD4 (no effect on DRD2) thereby reducing DRD4-dependent signal transduction at the post-synaptic membrane (loss of DRD4-dependent inhibition of adenylate cyclase (AC) resulting in an increase of cAMP production and PKA activation). Secondly, it promotes the expression of the DA-degrading enzymes COMT, MAOA and MAOB, thus exacerbating DA breakdown, globally reducing DA concentration at the pre-synaptic terminal and at the synaptic cleft. As a whole, ADAM10 inhibition, via the above described effects, would perturb the entire dopamine cycle and would ultimately lead to cognitive deficits (decrease in DRD4-mediated working memory) as well as behavioral impairments that could altogether contribute to the development of AD pathology.

In conclusion, our present study sheds new light on the mechanisms likely to induce an alteration of dopaminergic transmission during the development of AD and suggests ADAM10 as an element that could participate in this process. In a near future, it will first be of interest to correlate the herein observed effects on DRD4, COMT, MAOA and MAOB under GI254023X-mediated inhibition with an increase in the amyloidogenic processing of βAPP given that the modulation of ADAM10 activity is accompanied by changes in Aβ production [69]. Secondly, it will be necessary to determine whether the effects observed under GI254023X treatments are imputable to a decrease in sAPPα production or rely on some other ADAM10-dependent mechanisms involving the metabolism of one or more of its multiple other substrates [70]. This would undoubtedly help identifying the key mechanistic insights through which the inhibition of ADAM10 catalytic activity operates on the dopaminergic system.

## CRediT authorship contribution statement

**Subhamita Maitra:** Conceptualization, Investigation, Methodology, Data curation, Writing – original draft. **Bruno Vincent:** Conceptualization, Formal analysis, Supervision, Funding acquisition, Project administration, Writing – review & editing.

## Acknowledgements

We would like to thank Dr Narisorn Kitiyanant (Institute of Molecular Biosciences, Mahidol University, Thailand) for providing us with the SH-SY5Y cell line and Prof. Duncan R. Smith for his kind gift of GAPDH and PKA antibodies and MTT assay reagents. This work was supported by Mahidol University (NDFR 13/2564) and The Thailand Research Fund (BRG6180002). SM was supported by a Mahidol University postdoctoral research sponsorship.

